# Building a pathway for diversity in plant sciences in Argentina: highlighting the work of women scientists through virtual activities

**DOI:** 10.1101/2020.11.27.401661

**Authors:** Gabriela Alejandra Auge, María José de Leone, Rocío Deanna, Sonia Oliferuk, Pamela Anahí Ribone, Elina Welchen

## Abstract

Encouraging the participation of a diverse workforce in academia increases plurality as it broadens the range of skills, ways of thinking and experiences. Institutions and professional societies have been putting efforts on building plans that help make workplaces, conferences, education and extension programs more relatable to a highly diverse population. Argentina has an overall gender-balanced workforce in the sciences (~53% women/total), with an even higher representation in disciplines related to plant sciences. However, media outlets and national conferences related to genetics, botany, plant physiology, ecology and molecular biology, fail to reflect those numbers as the proportion of women invited for interviews, plenary lectures, and symposia falls below ~30%. As a way to increase the visibility of the wealth of plant science topics and experimental approaches in which Argentinian women work, and to facilitate connections among them across the country and abroad, we created the Argentinian Women in Plant Science network (https://argplantwomen.weebly.com/). This group has grown to over 200 members, representing a wide range of career stages and research topics. Since April, and taking advantage of the confinement situation, our weekly webinar series highlighting women plant scientists has reached an average audience of 60-70 participants, with a record of 100. Recently, we have begun a series of open professional development webinars to reach a wider public. Our first webinar, focused on Scientific poster design, had ~250 participants, most of them undergrad and graduate students from all over the country covering a diverse range of disciplines, including the social sciences. Even though we have immersed ourselves in the plant science community with our weekly seminars, we have expanded our goals with activities aimed to reach out to a much wider audience with webinars and teacher training workshops, hopefully making plant science more attainable to all.

## 1. Introduction

A diverse, gender-balanced environment improves working and learning experiences in the science workforce as a whole, increasing collective performance [1–4]. However, according to UNESCO, women account for a minority of the research workforce worldwide [5]. The numbers reflect gender disparity in scientific society leadership, conference speakers, journal editors and editorial board members, professorships, participation in media outlets, etc [6–9]. Many efforts have been made in recent years to overcome this problem and increase women representation in the sciences [10–16], but there is still a long road to break the ‘glass ceiling’ [17].

Argentina has an overall gender-balanced science workforce (~53%) [5], with even higher proportions of women in disciplines related to plant science (Figure 1). However, many science-related activities, especially in conferences and media outlets, fail to comply with this representation. With a group of Argentinian women plant scientists, we created the ARG Plant Women network to increase women visibility and interaction across the plant sciences in the country. Our objectives are to provide a network that facilitates connections and communication among its participants, to develop professional development opportunities, and to show the wealth of subdisciplines and experimental approaches used at work. In this short manuscript we would like to summarize gender representation numbers in our country, actions taken by the network and its initial outcomes.

**Figure 1.**
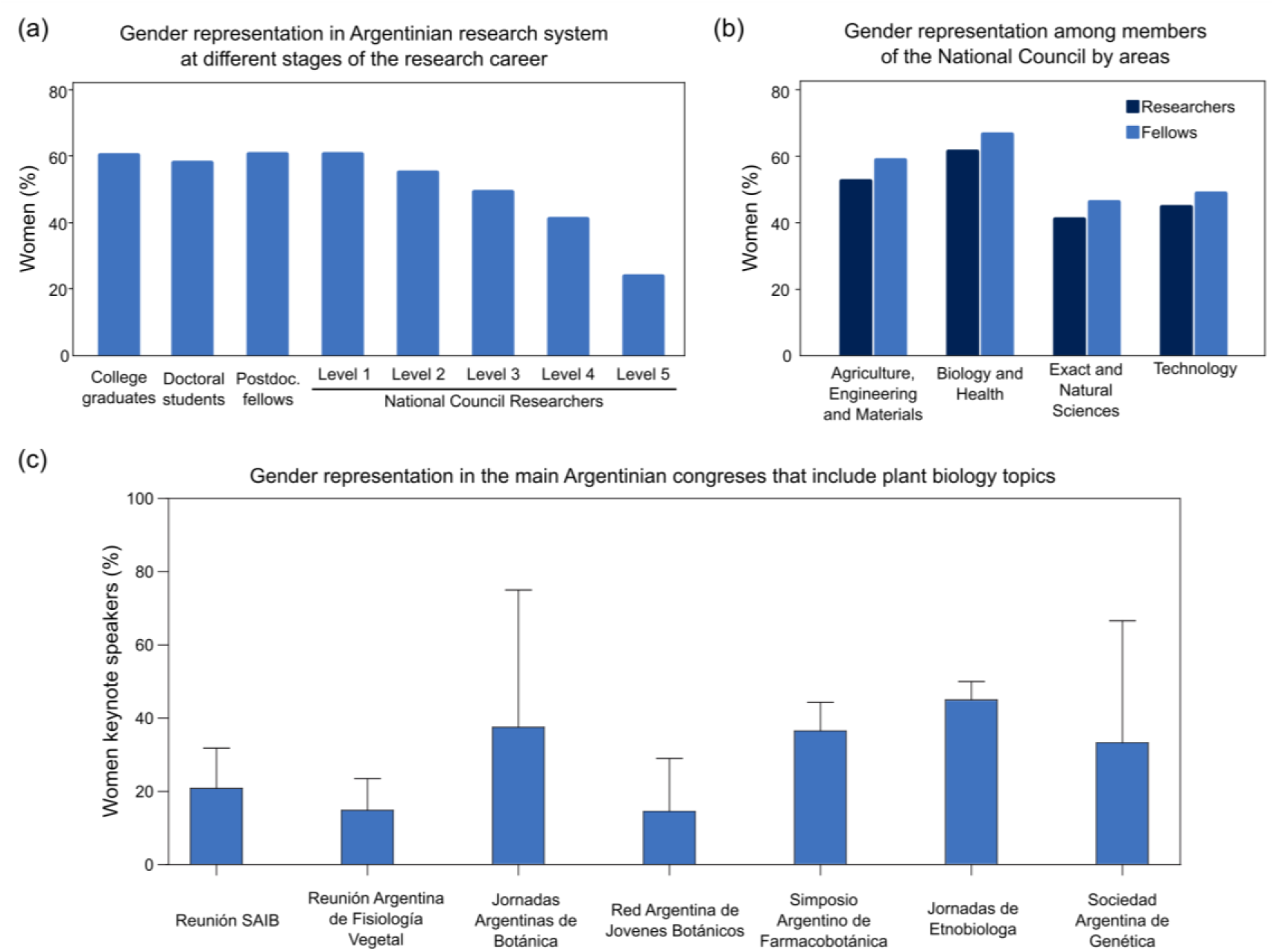
Gender representation of scientists in Argentina. **(a)** Proportion of women at different stages of the academic path (data: SICYTAR, Science and Technology Information Portal, and DIU, Department of University Information, Ministry of Education, Argentina [18,19]). **(b)** Proportion of women (National Council researchers, dark blue, and doctoral and postdoctoral fellows, light blue) in the main research areas that pertain to plant science (data: CONICET, National Council database). **(c)** Proportion of women invited as keynote speakers in national conferences. Data represent the average of the proportion of invited women scientists in the last 3 conferences of each area; ‘Reunión SAIB’, annual meeting of the Society for Research in Biochemistry and Molecular Biology; ‘Reunión Argentina de Fisiología Vegetal’, biannual meeting of the Society of Plant Physiology; ‘Jornadas Argentinas de Botánica’, biannual meeting of the Botanical Society; ‘Red Argentina de Jóvenes Botánicos’, biannual meeting of the Young Botanists Network; ‘Simposio Argentino de Farmacobotánica’, Pharmacobotany Symposium (every 4 years); ‘Jornadas de Etnobiología’, biannual Ethnobiology meeting; ‘Sociedad Argentina de Genética’, annual meeting of the Genetics Society.

## 2. Gender representation in the plant sciences in Argentina

Argentina is among the few countries in the world where gender parity in the sciences is reached [5]. Women represent most of the college graduates, doctoral students and postdoctoral researchers (Figure 1a). After a transition to an independent career as researchers of the National Council (Consejo Nacional de Investigaciones Científicas y Técnicas, CONICET), the most common path taken by academics in the country, we observe a sharp drop in the proportion of women as researchers are promoted to higher level positions (Figure 1a). This is a common trend in other positions related to education and science at universities and organizations in Argentina: fewer women in decision making positions (data not shown). Particularly in the major research areas of the National Council that include plant biology related topics, both women researchers and fellows are always well-represented (Figure 1b). However, we fail to see this gender inclusivity in speakers at national conferences related to genetics, botany, plant physiology, ecology and molecular biology, as the proportion of women invited for plenary lectures do not reflect those numbers, and falls below ~30% (Figure 1c).

## 3. ARG Plant Women: the Argentinian network of women in plant sciences

Even though plant sciences is a field with an overall good representation of women in our country, we observed that many seminars, conferences and activities fail to represent such numbers (Figure 1c). With the idea of increasing visibility of the many women that work in the plant sciences in the country, we created the network of Argentinian women in plant sciences, or ARG Plant Women [20]. Facing the lockdown due to the pandemic, we started thinking about ways to gather members of our network, and to create a virtual space we could use to know each other. The network now counts with over 200 members, with an extended community of ~250 people. The first activity we embarked on was a weekly virtual seminar series in which we invite women plant scientists, working in Argentina and abroad, to present their research and other science related activities (Figure 2a, Appendix A). From April to November, we have held 28 seminars with an average audience of 60 people, who stayed connected for most of the seminar (Figures 2b and 2c). Our speakers are scientists at different stages of their careers, and they work in many subdisciplines within plant science (Figure 2c). Appendix A shows a comprehensive list of the speakers, date of the seminars, affiliation and role, field of work and seminar title.

**Figure 2.**
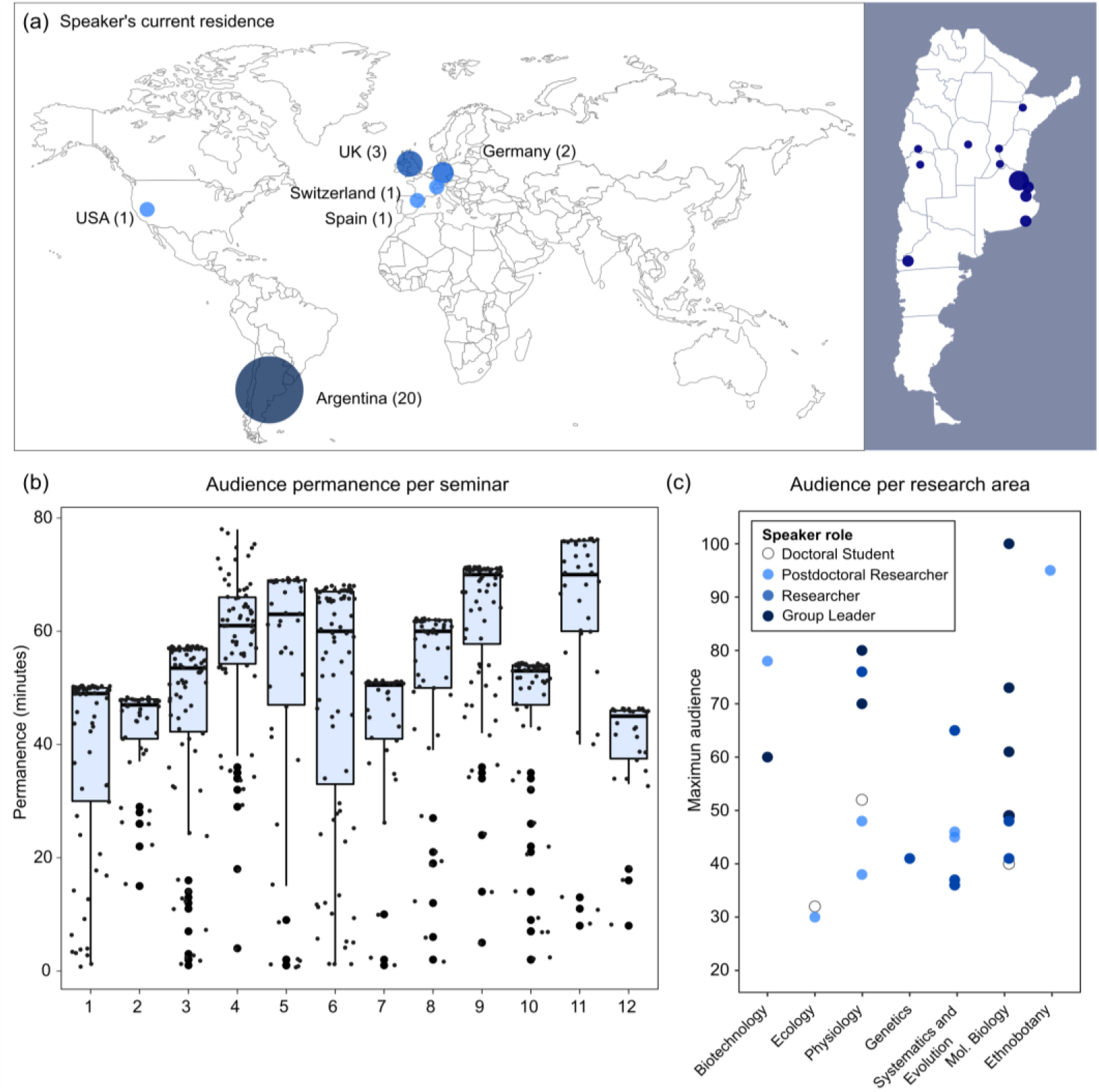
ARG Plant Women seminars. **(a)** Location and number of speakers per country in parenthesis. **(b)** Boxplots showing permanence in minutes of each person in the audience of the first 12 seminars (April-June 2020), note the median is close to the highest permanence points for all the seminars indicating most of the audience remains connected during the whole presentation. **(c)** Peak audience for each seminar (except the last 3 ones, y-axis) separated by subdiscipline in the plant sciences (x-axis) and career stage (color gradient).

We have also started a series of free open professional development webinars with the objectives of providing useful tools and reaching out to a wider public. The first two webinars (science poster design and effective presentations part I) counted with over 200 registrations each, an average of 100 active participants during the webinar and continuous participation in two dedicated Slack channels. The audience exceeded the plant science field as we had participants from the life sciences in general, but also some from the social sciences and even high school students. We observed great interest in the webinars and participation from undergraduate and postgraduate students, showing there is a need for this kind of courses that complement scientific training and personal development. We have scheduled two more webinars for before the end of the calendar year 2020 (effective presentations part II and effective communication skills). We have received many requests to repeat our webinars and to develop new workshops on other topics (public policy, student-mentor relationship, grant and fellowship applications, etc). In addition, and if the pandemic allows in-person conferences, we will join scientific meetings to offer a shorter version of these webinars, potentially increasing the audience reached.

## 3. Outcomes and self-evaluation

To measure the impact and level of acceptance of our seminar series, in July 2020 we conducted a survey among members of the network and seminar attendees. Our weekly virtual seminar series is highly rated among our network (Figure 3a). Half of the respondents considered the seminars changed their point of view about topics covered and the majority considered that change to be positive (Figure 3b). Many of the respondents have also taken actions as a result of attending the seminars, such as contacting speakers to explore new collaboration projects (Figure 3c). People highly valued different aspects of the seminars, like their format, the speaker selection and interaction with them, and also the availability of the recorded seminars on YouTube [21]. We have received an outstanding number of optimistic reviews and encouragement for the work we have done so far (Figure 3d). Nevertheless, we are still thinking about new ways to maintain the interest for the seminar series, increase diversity of themes and geographical representation, and at the same time to take into consideration the potential effects of the eventual re-opening of the workplaces for the next cycle 2021.

**Figure 3.**
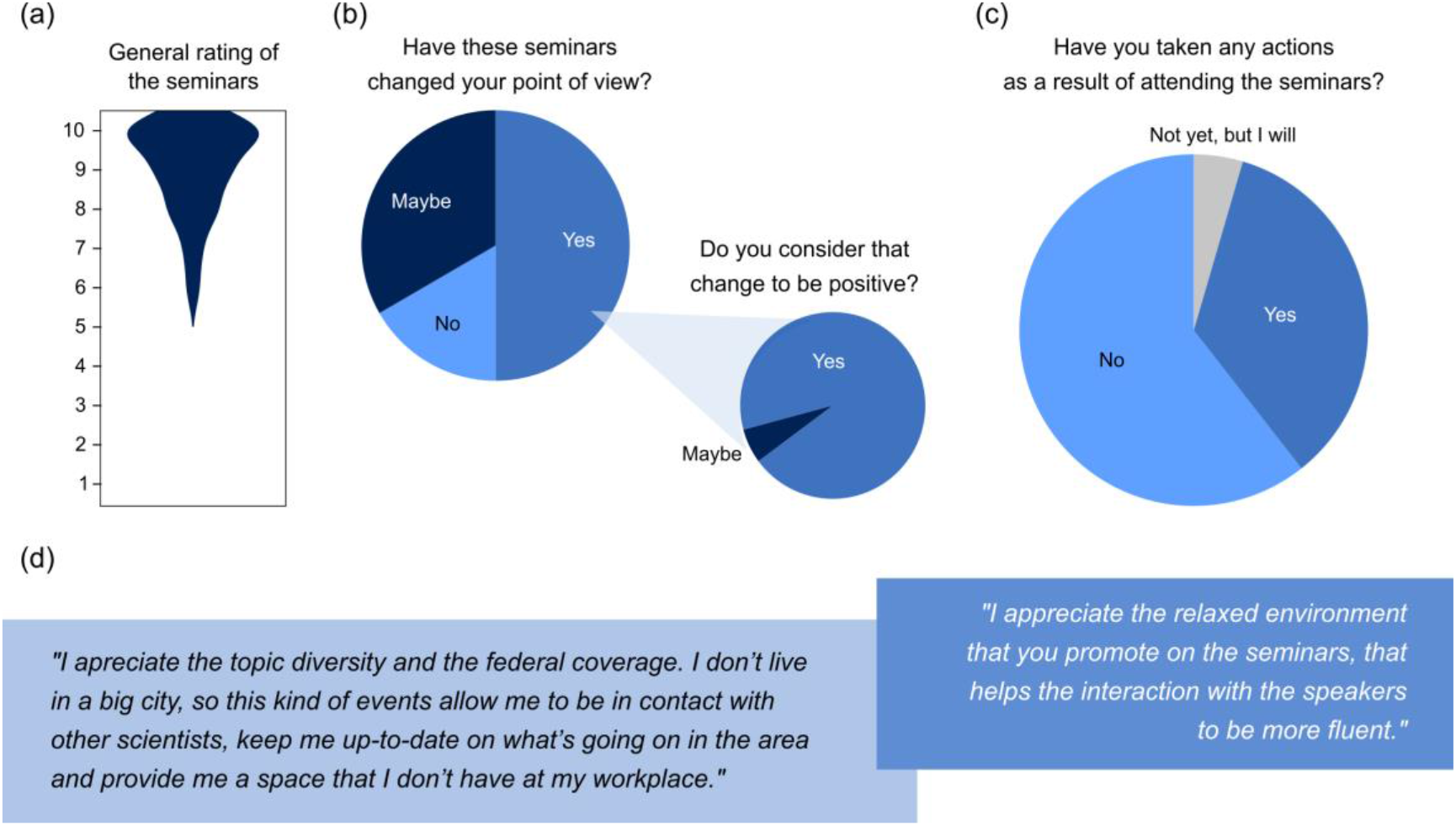
Seminars rating. **(a)** General rating of the seminars showing overall replies about quality. **(b)** Proportion of respondents that consider the seminars to effectively have changed (‘Yes’), might have changed (‘Maybe’) or have not changed (‘No’) their point of view; and for the ones that responded ‘Yes’, proportion of respondents who considered that change to be positive or not. **(c)** Proportion of respondents that have taken (‘yes’), will take (‘Not yet, but I will’) and have not taken actions (‘No’) after attending the seminars. **(d)** Exemplary comments left by the respondents.

On a similar survey, our webinars have received an average rate of 9.5 out of 10 points. The audience positively valued the duration, contents presented and format of the webinars, as well as interaction with and clarity of the speakers, extra material provided and proposed activities, plus communication through the Slack channels. In average, 97 % of the audience considered that they would apply the knowledge acquired in the webinars in their work. The large acceptance of these webinars shows we found a niche that needed to be covered for the career development of scientists, not only in our discipline, but in the whole Argentinian scientific community.

Engagement with our social platforms is active, with many followers and subscribers, as shown by metrics indicating interaction with our posts and publications (Table 1). The seminar videos, uploaded to the YouTube channel after the presentations, gathered over 1800 views.

**Table 1.** Engagement in social platforms: statistics from our Twitter, YouTube and Instagram accounts by November 2020.

| Social Platform | Subscribers /Followers | Impressions | Likes | Other metrics |
| --- | --- | --- | --- | --- |
| Twitter | 666 | 31.5K/month | ~340/month |  |
| YouTube [21] | 94 | 34.3K total | ~120 total | ~1800 total views |
| Instagram | 508 |  | ~280/post |  |

## 4. Next steps

Our aim is to make this network grow larger, reaching to many more places and scientists in our country we have not previously reached. We will keep working on increasing geographical representation and creating connections between speakers and audience, building stronger connections with Argentinian women plant scientists that are residing abroad, and developing more tools for personal development and knowledge of the intricate scientific system in our country. Even after re-opening of workplaces in 2021, we will keep working on a weekly seminar schedule in which graduate students, postdocs and researchers can show their work, as well as monthly webinars on different topics. We will develop teacher training workshops with the goal to incorporate more plant sciences in schools, hopefully encouraging more children and teenagers to pursue a career as plant biologists and botanists.

## 5. Conclusions

Even facing an ongoing pandemic, we have built a pathway to increase diversity in the plant sciences in Argentina. We believe there are many ways to promote the engagement and visibility of the work performed by women scientists, and we are delighted to see our work helps reach this goal. From this southern part of the world, we encourage to extend our planification to other countries and communities with the hope that this could be applied to further communication in science.

## Acknowledgments

G.A.A. is a researcher of the National Council (Consejo Nacional de Investigaciones Científicas y Técnicas, CONICET) and is supported by Agencia Nacional de Promoción de la Investigación, el Desarrollo Tecnológico y la Innovación (Agencia I+D+i, FONCyT, grants PICT 2016-0389 and PICT 2017-2656). M.J.dL. is supported by a CONICET fellowship. R.D. is supported by the National Science Foundation (DEB-1557871) and Agencia I+D+i (FONCyT, grant PICT 2017-2370). S. O. is supported by a fellowship from Agencia I+D+i (FONCyT, grant PICT 2016-0621). P.A.R. is supported by an EMBO long-term fellowship. E.W. is a researcher of the National Council (CONICET) and is supported by Agencia I+D+i (FONCyT, PICT 2017-2350). We would like to thank every member of the ARG Plant Women network and the plant science community at large for the continuous support to all our initiatives.

## Author Contributions

G.A.A., M.J.dL., R.D., S.O. and P.A.R. have gathered the data and wrote the manuscript. E.W. has contributed to the critical reading and final edition of the manuscript.

**Appendix A.** List of seminar speakers showing seminar date, affiliation, role field and seminar title. Videos available in our <u>YouTube Channel</u>.

| Date | Speaker | Affiliation | Role | Field | Topic |
| --- | --- | --- | --- | --- | --- |
| 6/4/2020 | Rocío Deanna | CU Boulder, US/Natural History Museum, UK | Postdoctoral Researcher | Systematics and Evolution | Evolución de la tribu *Physalideae* (Solanaceae) + Difusión de ciencia |
| 13/4/2020 | Anahí Yáñez | CIGEOBIO, CONICET-UNSJ, AR | Postdoctoral Researcher | Genetics | Retroelementos como marcadores moleculares en ají (*Capsicum*, Solanaceae) |
| 20/4/2020 | Mariana Corigliano | INTECH, CONICET-UNSAM, AR | Researcher | Biotechnology | HSP90 de plantas como inmunomoduladores de la RI + Divulgación y extensión |
| 27/4/2020 | Gabriela Amodeo | FCEN-UBA & IBBEA UBA-CONICET, AR | Group Leader | Physiology | Acuaporinas en plantas: los nuevos desafíos |
| 5/5/2020 | Laura Las Peñas | IMBIV, CONICET-UNC, AR | Researcher | Systematics and Evolution | Diversidad de *Cactaceae*: aspectos genéticos, morfológicos y ecológicos |
| 11/5/2020 | Gabriela Auge | IB3, FCEN-UBA, AR | Group Leader | Physiology | Vías de floración regulan pleiotrópicamente dormición y germinación de semillas |
| 18/5/2020 | Agostina Sassone | IPK - Leibniz Institute, DE | Postdoctoral Researcher | Systematics and Evolution | Dilucidando la taxonomía de los ajos sudamericanos (tribu *Leucocoryneae*) |
| 1/6/2020 | Gabriela Pagnussat | IIB, CONICET-UNMDP, AR | Group Leader | Molecular Biology | Síntesis de homocastasterona en el gametofito femenino de *A. thaliana* |
| 8/6/2020 | Martina Legris | Fankhauser's lab, UNIL, CH | Postdoctoral Researcher | Physiology | Rol de los fotorreceptores en el control del desarrollo en *Arabidopsis* |
| 16/6/2020 | Paula Casati | CEFOBI, CONICET-UNR, AR | Group Leader | Molecular Biology | Flavone biosynthesis and salicylic acid hydroxylation in plants |
| 25/6/2020 | Mariana Grossi | División Plantas Vasculares, Museo de La Plata, AR | Postdoctoral Researcher | Systematics and Evolution | Análisis comparativo de caracteres reproductivos en Eupatorieae (*Asteraceae*) |
| 29/6/2020 | Ana Srur | IANIGLA-CONICET, AR | Researcher | Ecology | Dinámica de los bosques en los Andes patagónicos argentinos: dendroecología |
| 3/8/2020 | Debora Rondanini | FAUBA, UBA, AR | Researcher | Physiology | Avances en ecofisiología de colza-canola |
| 10/8/2020 | Eugenia Zanetti | IBBM-FCE-UNLP, CONICET, AR | Group Leader | Molecular Biology | To keep or not to keep: mRNA stability and translatability in root nodule symbiosis |
| 24/8/2020 | Alicia Lopez Mendez | FCA UNMDP, CONICET, AR | Researcher | Systematics and Evolution | Nuestras líneas de investigación en botánica |
| 31/8/2020 | Julia Qüesta | CRAG, ESP | Group Leader | Molecular Biology | Epigenetic memory in plants |
| 7/9/2020 | Victoria Gastaldi | IAL, AR | PhD Student | Molecular Biology | Las proteínas TCPs y su participación en la elongación del estambre en *A. thaliana* |
| 14/9/2020 | María Laura Vidoz | IBONE, UNNE, CONICET, AR | Researcher | Physiology | Respuestas adaptativas de las plantas de tomate a la inundación |
| 21/9/2020 | Ana Ladio | INIBIOMA, AR | Researcher | Ethnobotany | Etnobotánica patagónica: vínculos con las plantas y su rol en conservación biocultural |
| 28/9/2020 | Pamela Ribone | SLCU, UK | Postdoctoral Researcher | Molecular Biology | Regulación de la expresión génica mediada por uORFs |
| 5/10/2020 | Luciana Bianchimano | FIL/IBBA-CONICET, AR | PhD Student | Physiology | Respuestas de las plantas al desafío impuesto por altas densidades de cultivo |
| 19/10/2020 | María Florencia Ercoli | Department of Plant Pathology, UC Davis, US | Postdoctoral Researcher | Molecular Biology | RaxX modifies the host environment by mimicking a plant peptide hormone |
| 26/10/2020 | Marina Clemente | INTECH, CONICET-UNSAM, AR | Group Leader | Biotechnology | Las plantas para la producción de proteínas de uso terapéutico + Ciencia en el bar |
| 2/11/2020 | Julietta Cagnacci | IIFA, INTA-CONICET, AR | PhD Student | Ecology | El impacto de diferentes ambientes naturales a lo largo de la altitud en las dinámicas de regeneración de dos especies de *Nothofagus* en los Andes Patagónicos |
| 9/11/2020 | Daniela Sueldo | SLS, Warwick University, UK | Group Leader | Molecular Biology | Plant immune cell death: execution and multicellular coordination |
| 16/11/2020 | Florencia Yannelli | Ecological Novelty group, FU-Berlin, DE | Researcher | Ecology | Restauración ecológica en el contexto de las plantas invasoras |
| 23/11/2020 | Rocío Tognacca | IFIBYNE, UBA-CONICET, AR | Postdoctoral Researcher | Molecular Biology | Regulación del splicing alternativo durante la germinación inducida por luz |
| 30/11/2020 | Agustina Yañez | MACN-CONICET, AR | Researcher | Systematics and Evolution | Biodiversidad de Helechos y Licofitas en áreas protegidas de la provincia de Misiones: 10 años de trabajo en la selva Paranaense |

